# Laboratory evolution reveals transcriptional mechanisms underlying thermal adaptation of *Escherichia coli*

**DOI:** 10.1101/2024.02.22.581624

**Authors:** Kevin Rychel, Ke Chen, Edward A. Catoiu, Connor A. Olson, Troy E. Sandberg, Ye Gao, Sibei Xu, Ying Hefner, Richard Szubin, Arjun Patel, Adam M. Feist, Bernhard O. Palsson

**Author notes:** **Corresponding author:** Bernhard O. Palsson.

## Abstract

Adaptive laboratory evolution (ALE) is able to generate microbial strains which exhibit extreme phenotypes, revealing fundamental biological adaptation mechanisms. Here, we use ALE to evolve *Escherichia coli* strains that grow at temperatures as high as 45.3°C, a temperature lethal to wild type cells. The strains adopted a hypermutator phenotype and employed multiple systems-level adaptations that made global analysis of the DNA mutations difficult. Given the challenge at the genomic level, we were motivated to uncover high temperature tolerance adaptation mechanisms at the transcriptomic level. We employed independently modulated gene set (iModulon) analysis to reveal five transcriptional mechanisms underlying growth at high temperatures. These mechanisms were connected to acquired mutations, changes in transcriptome composition, sensory inputs, phenotypes, and protein structures. They are: (i) downregulation of general stress responses while upregulating the specific heat stress responses; (ii) upregulation of flagellar basal bodies without upregulating motility, and upregulation fimbriae; (iii) shift toward anaerobic metabolism, (iv) shift in regulation of iron uptake away from siderophore production, and (v) upregulation of *yjfIJKL*, a novel heat tolerance operon which we characterized using AlphaFold. iModulons associated with these five mechanisms explain nearly half of all variance in the gene expression in the adapted strains. These thermotolerance strategies reveal that optimal coordination of known stress responses and metabolism can be achieved with a small number of regulatory mutations, and may suggest a new role for large protein export systems. ALE with transcriptomic characterization is a productive approach for elucidating and interpreting adaptation to otherwise lethal stresses.

## Introduction

Adaptive laboratory evolution (ALE) can generate microbes which push biological systems to extremes, which enables the study of interesting new phenotypes and can provide insight into fundamental biology. Starting with a microbe and condition of interest, cells are grown for many generations and propagated in exponential growth phase when flasks reach a target density (1). Mutants that spontaneously arise in the flask and are able to grow faster under the given condition will have a higher likelihood of being propagated, so the population will accumulate beneficial mutations and evolve. For tolerization ALEs, cells are grown in a stressful condition, and the stress is increased when the flask’s growth rate stabilizes, enabling the development of highly stress-tolerant strains (2). A detailed understanding of these strains reveals mechanisms of stress tolerance which are able to inform the design of cellular factories (2), our understanding of the evolution of pathogens (3), and the fundamental science of systems that may otherwise be hard to study.

Typically, ALE endpoint strains are studied by DNA resequencing and subsequent characterization of the mutations that are expected to improve fitness. This works well for many ALEs, as evolved strains typically have a relatively small number of mutations (4). However, microbes are able to increase their mutation rate by mutating the DNA mismatch repair machinery, and will evolve into hypermutator strains in highly stressful environments (5). These strains are therefore of particular interest, since they rapidly acquire novel phenotypes. However, it is very difficult to elucidate the key causal mutations from hypermutator strains due to the high number of mutations, each of which may interact with one another in complex ways or have very little effect at all. Therefore, methods that can reduce the complexity of analyzing hypermutator strains and cut through the genomic noise of their many mutations are needed.

The transcriptional regulatory network (TRN) senses cellular states and environments, and helps to maintain homeostasis and regulate growth by adjusting gene expression levels. Analyzing changes to transcriptomic allocation in hypermutator strains represents a possible route for their characterization, as it can reveal regulatory changes induced by the mutations or be used to infer changes to metabolism and stress. However, we again run into a problem of scale: hundreds or thousands of genes may be differentially expressed, making it difficult to glean a global understanding from transcriptomic datasets.

This problem can now be addressed by a transcriptomic analysis approach called independently modulated gene set (iModulon) analysis (6). iModulon analysis employs independent component analysis (ICA) to identify co-regulated signals from large compendia of gene expression data. These signals are represented by iModulons, which have a weighting for each gene in a signal and an activity level (signal strength) in each sample. Highly weighted genes are considered to be members of the iModulon. iModulon gene sets tend to match well with regulons. Regulons are defined using a variety of bottom-up experimental methods (7) whereas iModulons are quantitative structures learned top-down from expression data alone. Compared to analyzing individual gene expression levels, analyzing the activity levels of iModulons decreases the number of significant variables approximately 17-fold (8). This simplification makes it tractable to characterize most of the variance in gene expression and interpret it using associations between iModulons and known transcriptional regulators. Since these signals are typically identified based on hundreds of transcriptomes, they also provide useful context for analyzing trends in new sets of samples. iModulon structures have been established for several organisms and are available to browse, search, or download from iModulonDB.org (9).

iModulons have proven useful for analyzing non-hypermutator ALE strains in several cases (10–15), making them a promising option for characterizing hypermutator strains. We recently compiled over 1,000 *E. coli* K-12 RNA sequencing (RNAseq) expression profiles into a dataset called PRECISE-1K and characterized its iModulons (8), which are available on iModulonDB.org. Data from the present work was included in that dataset, but presented without the detailed characterization provided here.

To explore how transcriptomic allocation adaptively evolves, we must choose a selection pressure that will produce informative strains. High temperature exerts a fundamental stress on biological systems by destabilizing proteins, membranes, and other molecules (16). Tolerating this stress has driven evolution since life began (17), is relevant to understanding the response of pathogens to fever (18), could be helpful to engineer more efficient cell factories (19), and will become a more common selection pressure as global warming continues (20). One prior study demonstrated the presence of epistasis among mutations acquired under this stress (21). Another prior study used ALE with increasing temperatures to generate ten *E. coli* strains that grow well at 42°C (22). Mutational analysis and a simple transcriptomic analysis of these strains revealed some valuable thermal adaptation strategies, such as modifying mRNA degradation and peptidoglycan recycling pathways. However, interpreting the transcriptomic responses of these strains was difficult in the absence of iModulon analysis, and we predicted that higher heat tolerance could be achieved with further evolution.

Here, we evolved an isolate from the 42°C evolution further to push the limits of heat tolerance. The six endpoint strains of this study can grow at temperatures as high as 45.3°C, which is lethal to wild type strains. To achieve this increase in heat tolerance, the strains were all hypermutators. We generated transcriptomic samples from these strains at various temperatures and previously included them in PRECISE-1K (8). We enumerated each of the major transcriptomic adaptations that facilitate rapid growth at high temperatures. Despite their broad range of genomic mutations, the strains exhibited only a few major transcriptomic changes. These correspond to the regulation of stress responses, motility, redox metabolism, and iron uptake. We also identify and predict protein structures for a strongly upregulated, previously uncharacterized operon, *yjfIJKL*, which may be beneficial for survival at high temperatures. In addition to the specific insights on heat tolerance, this study demonstrates the value of transcriptomic analysis (particularly the iModulon framework) for gaining clear insights from hypermutator strains.

## Results

### ALE increased temperature tolerance via a hypermutator phenotype

Using ALE, we obtained six evolved strains that tolerated 45.3°C (**Figure 1A, S1**). Each one descended from the same ancestor from the previous 42°C ALE (22), 42c_3, which itself descended from *E. coli* K-12 MG1655. The mutations present in the starting strain are summarized in **Table S1.** We generated growth curves and computed growth rates for each of the strains at 30, 37, and 44°C (**Figure 1B**), showing a significant increase in growth rate at 44°C after evolution (p = 5.9*10^-7^). Interestingly, the strains did not exhibit a major tradeoff in growth rates at 37°C on average (p = 0.13), and only had a slight growth disadvantage against wild type at 30°C (p = 0.012). The evolved strains maintained much of their ability to adapt to changes in temperature, suggesting that the acquired genomic mutations were not especially deleterious at lower temperatures.

**Figure 1.**
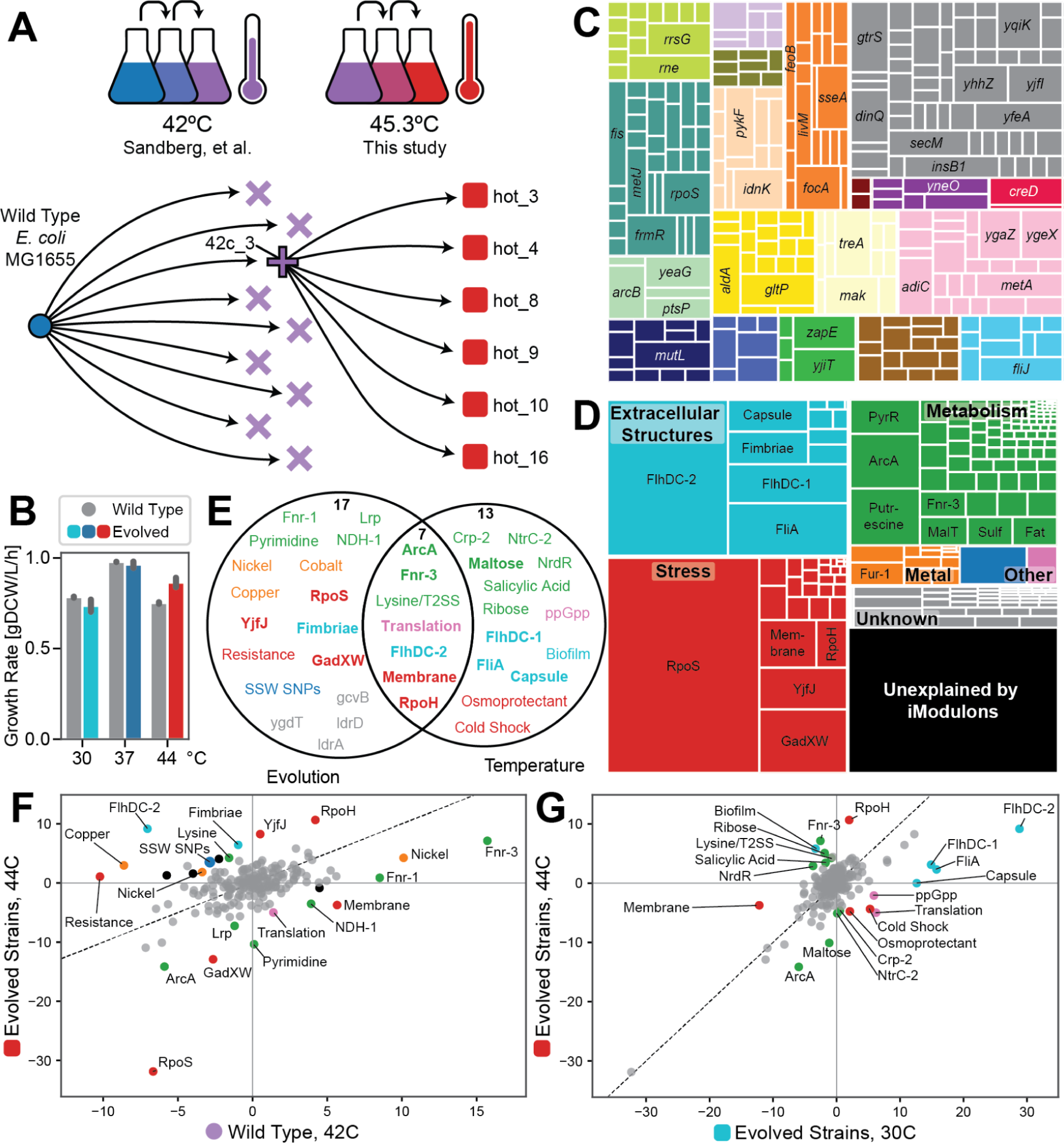
ALE increased temperature tolerance via changes to the genome and transcriptome. **(A)** ALE schematic, showing a previous round of ALE that tolerized *E. coli* up to 42°C (22), and that the present study focuses on descendents from a single strain of the prior study to generate strains that tolerate up to 45.3°C. Symbol shapes represent strain cohorts, and colors represent temperatures; these will be kept consistent throughout the paper. See **Figure S1** for details. **(B)** Growth rates for the wild type and final six evolved strains at three temperatures, showing a significant increase in growth rate at 44°C (p = 5.9*10^-7^) but similar growth rates as WT at 30 and 37°C. **(C)** Treemap of all 504 mutations observed in any of the six evolved strains, where each mutation is mapped to its nearest gene and genes that mutate in five or more strains are labeled. Colors indicate clusters of orthologous genes (COGs) (**Table S2**). **(D)** Treemap of the variance in the transcriptomes of the evolved strains by iModulon, showing that relatively few iModulons capture most of the variation. The 20 iModulons which explain the most variance are labeled, with some names shortened for space. Categories are labeled with bold names, except two categories combined as “Other”: “Genomic” in blue and “Translation” in pink. For more information on each iModulon, see **Table S3** and iModulonDB.org (*E. coli* PRECISE-1K). **(E)** Venn diagram of the 37 significant differential iModulon activities (DiMAs) from evolution (F) and temperature changes (G). Colors match the categories in (D). Bold iModulon names are also in the top 20 by explained variance. **(F-G)** Colors match the categories in (D), except that gray represents insignificant iModulon activities and black represents the “unknown” category. **(F)** DiMA plot comparing the iModulon activities in the wild type and evolved strains at their highest respective temperatures. **(G)** DiMA plot comparing iModulon activities in the evolved strains at cold (30°C) and hot (44°C) temperatures.

The ancestral strain, 42c_3, contained 30 mutations, including *mutL* G49V. MutL is part of the DNA mismatch repair machinery, and mutations in this gene tend to induce increases in mutation rates (5, 23–25). Thus, each of the evolved strains was a hypermutator, and ended with between 60 and 126 mutations, with the average strain experiencing 84 mutations. Details of all mutations are shown in **Table S2**. Each of the mutations was assigned to its nearest gene, and mutated genes were visualized in **Figure 1C**. No particular cluster of orthologous genes (COG) was enriched in this set, and the large number of mutations precluded a detailed analysis of the potential benefit of each one, as has been performed in past ALE studies (e.g. (22)).

### iModulon analysis revealed transcriptomic adaptations

In order to gain a clear understanding of the adaptations in the evolved strains, we generated RNAseq data for each of them and the 42c_3 ancestor at 30, 37, and 44°C in duplicate. From the prior study (22), we also had samples at 42°C for the wild type and each of the 42°C evolved strains. All of these profiles were included in PRECISE-1K, a compendium of over 1,000 *E. coli* transcriptomes which were generated using the same experimental protocol and analyzed with iModulons in aggregate (8).

PRECISE-1K provides a large and diverse condition space from which to identify co-regulated, independently modulated signals (iModulons). The 201 iModulons computed from PRECISE-1K have been characterized with assigned functions, regulators, and categories to facilitate interpretation. They are available at iModulonDB.org (9) under “*E. coli* PRECISE-1K”, in the project “hot_tale”. We can quantify the explained variance of each iModulon in the samples from the evolved strains (**Figure 1D**). In the data generated for this study (a subset of PRECISE-1K), the 201 iModulons captured 82% of the variance in the data, with the remaining 18% assumed to be noise or other variation with no clear structure. The top 20 highest explained variance iModulons explained 61.3% of the overall variance (labeled iModulons, **Figure 1D**). Thus, a relatively small number of variables can be highly informative about the global state of these samples, and therefore represent an approach to identify key thermotolerance strategies that emerge during ALE.

Differential iModulon activities (DiMAs) are similar to commonly used plots of differentially expressed genes (DEGs), except they are much easier to interpret because the iModulons are much fewer in total number (201) than genes (4257). iModulons are also knowledge-enriched with regulatory information (8). DiMAs between starting and evolved strains represent a summary of transcriptional adaptations (14) (**Figure 1F**). In this dataset, we can also quantify DiMAs for the evolved strains between the cold and hot temperatures (**Figure 1G**), and compare the sets of significant iModulons (**Figure 1E**). Statistics and additional details for each iModulon are provided in **Table S3**.

Highly variable and differentially activated iModulons in ALE studies typically indicate one of three features (14): (1) large genomic alterations have directly amplified or deleted the genes of an iModulon, (2) mutations in regulatory pathways have altered gene expression, or (3) underlying metabolites or processes which are sensed by the TRN have been altered. There were no major amplifications or deletions in the evolved strains which resulted in their own iModulons. Therefore, the major signals are either the result of regulatory mutations or the output of the sensory systems within the evolved cells.

Based on the direction of the change, the large body of existing literature on the *E. coli* TRN, and experimental evidence, we have inferred the mechanisms which underlie the major changes in the transcriptome. We also proposed explanations of how they provide benefits to the evolving strains. We present the five mechanisms with the strongest signals in the following sections, in **Figure 2**, and in **Table S4**.

**Figure 2.**
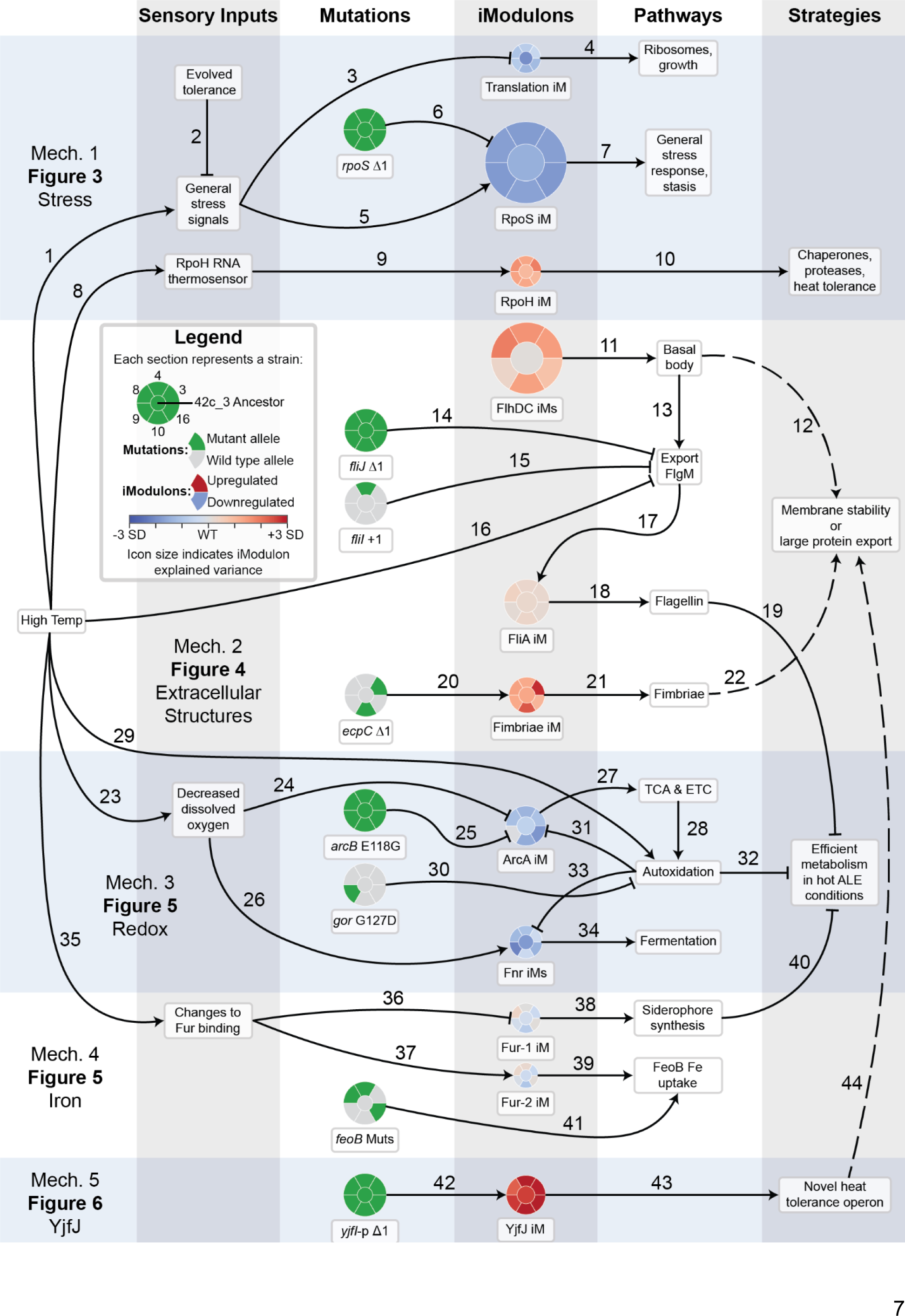
Overview of the five major mechanisms (mechs) that underlie the adaptation to high temperature growth. Each relationship marked by an arrow corresponds to a row in **Table S4**, in which its evidence and novelty is listed. On the left side, high temperature is the main “input”, which leads to a variety of sensory inputs in the first column, which are expected from literature or inferred from the iModulon evidence. In the second column, presence or absence of mutations in the evolved strains are shown, according to the legend. iModulons in the third column integrate sensory inputs and effects of mutations to determine their activity levels. Colors in the iModulon icons represent the change between the wild type (WT) activity at 42°C and the given strain’s evolved activity at 44°C, normalized by the standard deviation (SD) of the iModulon’s activity in all samples from PRECISE-1K (e.g. the RpoS iModulon is downregulated (blue), the FliA iModulon is constant (gray), and the YjfJ iModulon is upregulated (red)). iModulon icons are sized according to their explained variance (from Fur-2, 0.27% to RpoS, 19.8%; scaled using the square root). To the right of the iModulon column are the pathways and phenotypes which are determined by the transcriptomic and genomic changes. The far right column lists hypothesized strategies by which the evolved strains tolerated high temperatures. Different background shades represent different topics/mechanisms, and each is labeled with the respective figure in which to find more information.

### Stress sigma factors shift transcriptomic allocation from general to specific responses

The iModulon with the largest explained variance in observed transcriptomic changes is the RpoS iModulon, which reallocates an enormous 20% of the transcriptome in the evolved strains. RpoS is the general stress response sigma factor, which is governed by complex regulation and limits growth when active (26, 27). Prior iModulon studies have explored a “fear/greed tradeoff”, where typical strains exhibit a negative correlation between the RpoS and Translation iModulon activities. Faster growing cells activate the Translation and downregulate the RpoS iModulons (6, 11, 14, 15, 28, 29). As the prior generation of 42°C evolved strains mutated to tolerate high temperatures, they experienced less stress and therefore downregulated RpoS (22) (**Figure 3A**, “42c Other”).

**Figure 3.**
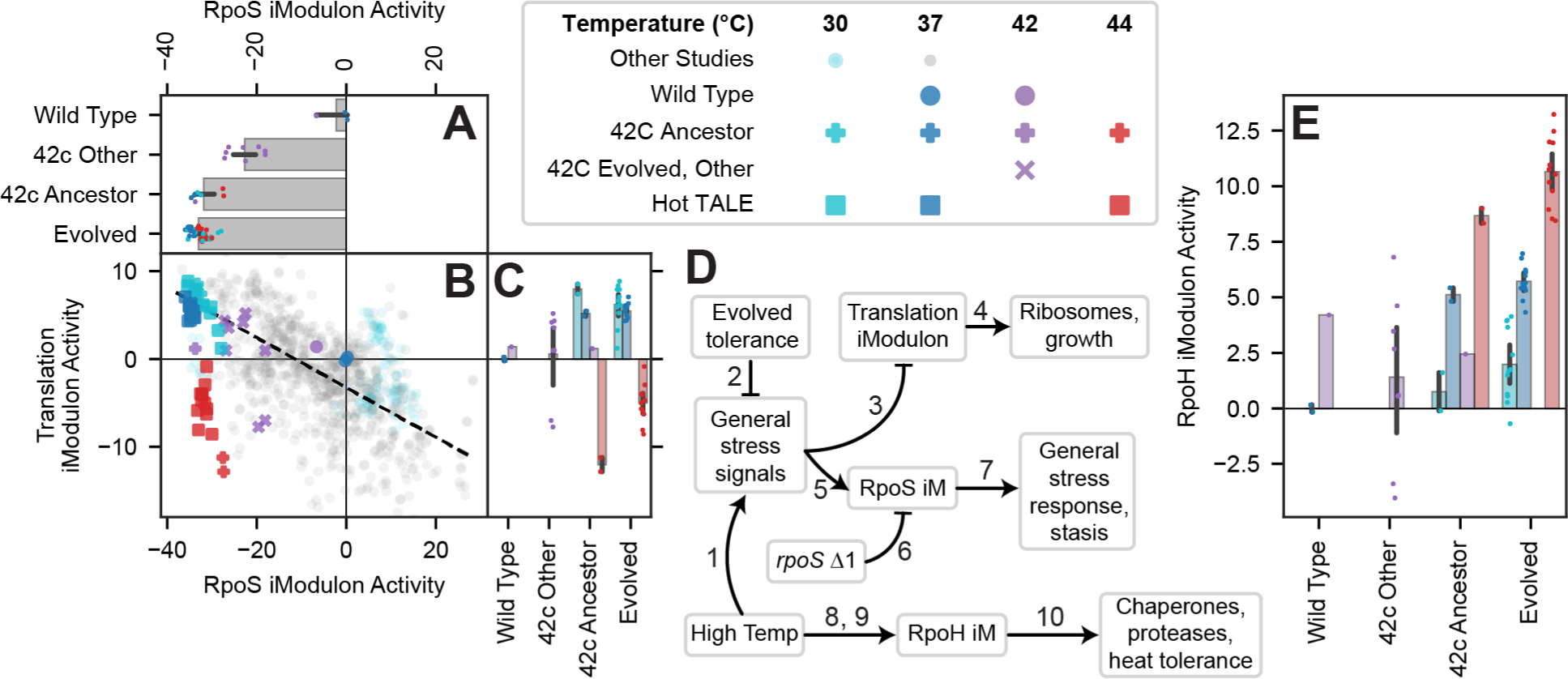
Stress sigma factors shift allocation from general to specific responses. Error bars represent mean ± 95% confidence interval. Colors in the columns of the legend are consistent for each plot. **(A-C)** Activities of the RpoS and Translation iModulons, which constitute the fear/greed tradeoff. **(A)** RpoS activity is downregulated by evolution (p = 0.036), and accounts for 19.8% of the variance in the dataset. The RpoS iModulon regulates the general stress response which slows growth. **(B)** Scatter plot color coded according to the legend, with low opacity circles representing all other samples in PRECISE-1K (n = 969). A black dashed line was fit to the other samples, representing the typical fear/greed tradeoff (Dalldorf et al. 2023). The temperature evolved samples have lower RpoS activity than expected, due to the *rpoS* mutation in the strains. **(C)** Translation activity is correlated with temperature, but downregulated less strongly after evolution due to successful adaptation (p = 0.027). **(D)** Knowledge graph describing this figure. Numbered arrows are consistent with Figure 2 and reference additional details in **Table S4**. **(E)** RpoH iModulon activity, which maintains its correlation with temperature but is slightly upregulated by prolonged heat exposure (p = 0.0031). RpoH regulates high temperature responses.

In the 42c_3 ancestor of the high temperature tolerant strains, we observe even stronger downregulation of the RpoS iModulon, which is maintained in the new evolved strains. This iModulon activity is likely resulting from a frameshift mutation in *rpoS* in 42c_3 and its derivatives. The mutation appears to have mostly deactivated RpoS, allowing sigma factors that do not suppress growth to outcompete it and providing a growth rate benefit during ALE. Deactivating RpoS is generally a good strategy for ALE (6, 11, 14, 28–31), but the temperature adapted strains take this to an extreme via this mutation.

The Translation iModulon is typically anti-correlated with RpoS, because similar underlying growth/stress and RNA polymerase-related variables control both iModulons (26, 27). However, the Translation iModulon remains anti-correlated with temperature (**Figure 3C**) while the RpoS iModulon is downregulated at all temperatures, diverging from the usual fear/greed tradeoff (**Figure 3B**). The *rpoS* frameshift stops RpoS activity but does not regulate the Translation iModulon, explaining this discrepancy (**Figure 3D**). We do observe a small upregulation of the Translation iModulon at high temperatures after evolution, suggesting that the mutations and tolerization strategies in these strains have successfully decreased the stress signals which typically downregulate Translation at high temperatures.

Unlike RpoS, the heat stress sigma factor RpoH does not mutate, maintains its wild type correlation with temperature, and is differentially upregulated at high temperatures after evolution (**Figure 3D**). RpoH senses temperature via several mechanisms including an RNA-thermosensor and temperature-dependent proteolysis (32, 33), and it activates a variety of heat shock genes and chaperones (34). Presumably, any mutations or changes to heat stress regulation were selected against. Prolonged exposure to high temperatures slightly upregulates RpoH, probably via the known temperature-dependent pathways (32, 33).

Thus, the evolved cells downregulate general stress responses (RpoS) to improve growth, but upregulate specific responses to heat (RpoH). This represents an effective strategy for stress tolerization ALE; indeed, it mirrors the response of oxidative stress evolved strains, which maintain activity of the specific oxidative stress response, SoxS, while also downregulating RpoS (14).

### Motility iModulons amplify flagellar basal body expression while suppressing the filament

A major fraction (18%) of the variance in the transcriptome of the thermotolerant strains is explained by motility iModulons, which respond to two transcription factors, FlhDC and FliA. The regulation of this system has been studied in detail (35), and the iModulon gene structure matches well with the known literature. The two primary iModulons of interest are FlhDC-2 and FliA (**Figure 4A**). The promoter of *flhDC* integrates many signals that affect motility (36–41), and then expression of FlhDC induces flagellar synthesis in steps: first, the basal body is synthesized, and then the hook, junction, flagellin, motor, and control mechanisms are added. The timing of these steps is ensured by using a second regulator, the sigma factor FliA, which is induced by FlhDC (as part of the class I genes, purple in **Figure 4A**), co-regulates the intermediate steps (class II genes, orange), and solely regulates the final steps (class III genes, green) (35). A good understanding of the regulation and dynamics of this system is important for fundamental biology, understanding host-pathogen interactions (42), and developing a toolkit for designing protein secretion systems (43, 44). For more information on these iModulons and FlhDC-1, see **Note S1**.

**Figure 4.**
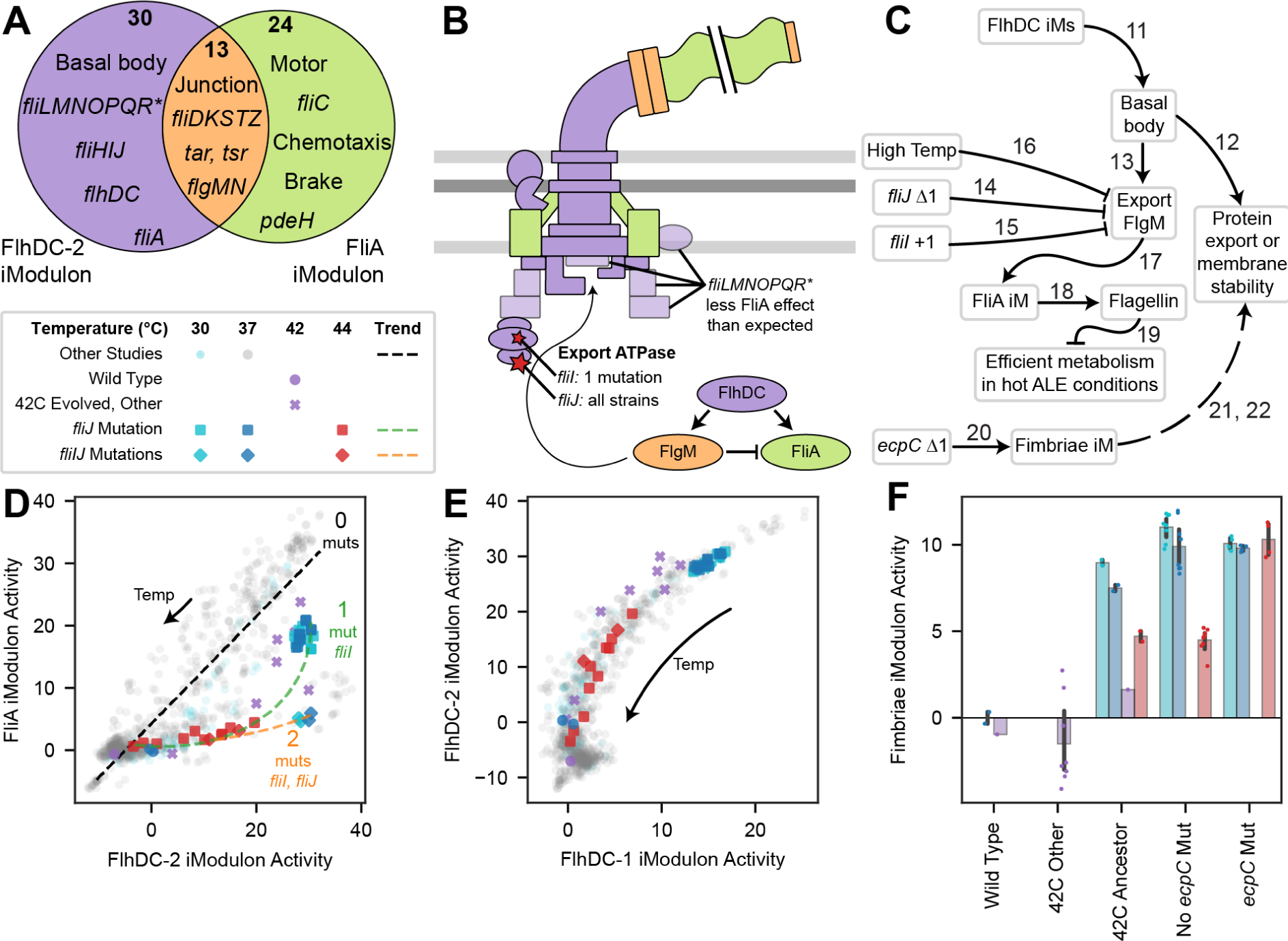
Changes to motility and fimbriae regulation suggest a possible role in high temperature tolerance. **(A)** Venn diagram of genes in the two main motility iModulons, FlhDC-2 and FliA, which are highly similar to the known regulons (Fitzgerald et al. 2014; Santos-Zavaleta et al. 2019). The most notable exception is *fliMNOPQR*, which is thought to be regulated by both FlhDC and FliA but is only found in the FlhDC related iModulon (**Note S1**). **(B)** Illustration of the flagellum, adapted from (35). Components are colored according to the Venn diagram in (A), and low opacity components are those which were expected to be under dual regulation based on the literature. ATPase mutations in *fliI* and *fliJ* observed in the evolved strains are pictured with a red star and label. The established regulatory cascade (35) is also pictured in the lower right, showing that the anti-sigma factor FlgM requires the ATPase to be exported and derepresses FliA. **(C)** Knowledge graph summarizing this section. Numbered arrows are consistent with Figure 2 and reference additional details in **Table S4**. **(D-E)** Scatter plots of iModulon activities (iModulon activity phase planes), illustrated according to the legend above panel (D). **(D)** Though FliA activity is typically correlated with FlhDC-2 activity (black dotted line: best fit for other projects; Pearson R = 0.87) and follows it in the regulatory cascade (B), the mutant strains (squares) do not activate FliA as strongly as expected, particularly in the case of the strain with two ATPase mutations (diamonds). Green and orange lines illustrate observed trends. Temperature and other regulatory changes move strains along the line, while mutations modify the line. **(E)** The activity of the two FlhDC iModulons form a tight curve with heat tolerant strains having high activity below 40°C but decreasing at high temperatures. This trend is not affected by observed mutations. **(F)** Bar and swarm plot of Fimbriae iModulon activity, with all evolved strains upregulating it (p = 0.00030) and those with the *ecpC* Δ1 mutation upregulating it the strongest at high temperatures. Error bars represent mean ± 95% confidence interval.

At high temperatures, the flagellar secretion system is less able to secrete FlgM, the anti-sigma factor for FliA (45). This mechanism may have evolved to help *E. coli* avoid flagellin-mediated detection by the host immune system during a fever (42). Thus, the flagella synthesis pathway is cut off at a regulatory point between the two iModulons (**Figure 4C**). We observe this mechanism clearly in the activity of the evolved strains at 44°C, which have some FlhDC-2 activity but no FliA activation (**Figure 4D**). The activity phase plane between FlhDC-2 and FliA is thus highly informative: typically, the two iModulons exhibit a strong correlation (Pearson R = 0.87), but in cases below the best fit line, a mechanism such as the failed secretion of FlgM inhibits FliA activity.

Interestingly, we also observed the evolved samples exhibiting activity below the best fit line at lower temperatures, when heat should not be disrupting FlgM secretion (**Figure 4D**). This observation indicates that something besides temperature in these strains downregulates the FliA iModulon. Indeed, the ancestor (42c_3) of all evolved strains had a frameshift mutation in *fliJ* (**Table S1**), which is involved in the secretion of flagellar export substrates (46). The strains occupy a unique location in the phase plane, possibly because FlgM can’t be exported as efficiently due to this mutation. Another mutation, a frameshift in the export ATPase *fliI*, affected only the hot_4 strain. With this mutation, FliA activity decreases even further (diamonds, **Figure 4D**).

iModulons have thus quantitatively captured the complex transcriptional regulation of motility and revealed the effects of temperature and mutations on the regulation of FliA. Key questions remain to be elucidated: (i) the strains strongly upregulate FlhDC, but due to its complex upstream regulation it is difficult to deduce the molecular mechanism, (ii) since FlhDC activity also decreases at high temperatures, some unknown mechanism downstream of FliA may be feeding back to regulate these genes, and (iii) there may be an evolutionary benefit to expressing FlhDC-regulated genes but not FliA-regulated genes at high temperatures.

We can speculate about the evolutionary benefit in question (iii), and propose two hypotheses which warrant further study: (a) the basal body could secrete other temperature-sensitive proteins, and/or (b) the basal body may provide membrane stability. (a) The flagellar basal body is able to rapidly secrete large proteins, and can be engineered to secrete a variety of protein substrates (43). Its typical secretion substrate, *fliC*, is downregulated by the evolved *fliIJ* mutants, which means that the exporter is upregulated while its substrate is absent. Therefore, perhaps it has been repurposed to help eliminate protein aggregates in the cells, which would accumulate under high temperatures. Alternatively, (b) the flagellar basal body is one of very few protein complexes that span both membranes. More basal bodies may therefore be structurally beneficial to the cellular envelope, which is destabilized by high temperatures. Both of these hypotheses warrant future study, and are also applicable to the Fimbriae and YjfJ iModulons discussed later.

### The Fimbriae iModulon is another upregulated large protein export system

Another extracellular structure, the fimbriae, is an important part of transcriptome reallocation in the evolved strains (1.6% explained variance). This iModulon contains the fimbriae synthesis genes *fimAICDFGH (47)*, which are strongly upregulated in 42c_3 and the evolved strains and negatively correlated with temperature **(Figure 4F)** (48). Interestingly, the negative correlation with temperature was abolished and fimbriae were strongly upregulated at high temperatures in two evolved strains, hot_3 and hot_10, which both shared a frameshift mutation in the *ecpC* gene. EcpC is a putative usher protein for another extracellular structure, the common pilus (49). This result suggests cross-talk between various extracellular fiber systems in *E. coli*, and it also suggests that upregulation of fimbriae may be beneficial at high temperatures.

Similar to our two hypotheses about flagellar basal bodies, the upregulation of fimbriae may also assist with misfolded protein export and/or membrane stability. The usher proteins *fimD* and *ecpC* typically export large proteins, and may also have been repurposed for misfolded substrates in the evolved strains. Unlike flagellar basal bodies, the fimbriae systems are only in the outer membrane and would therefore be limited to secreting periplasmic proteins or supporting the stability of the outer membrane. Further research into the specificity of protein export by fimbriae and pilus systems, and their role in thermal stability of the cellular envelope, could be helpful for understanding heat tolerance and for the design of heterologous protein producers (50).

### Redox metabolism shifts toward less aerobic metabolism

ArcA and Fnr, the regulators of aerobicity, exert significant control over cellular phenotypes via alterations to the expression of genes involved in respiration (51). Together, their associated iModulons explain 3.4% of the variance in the transcriptomes of the evolved strains, but are likely to have a larger effect on metabolism and phenotypes. The ArcAB two-component system represses aerobic metabolism genes when the electron transport chain (ETC) is in a reduced state (52), and Fnr derepresses anaerobic metabolism genes when its iron-sulfur (Fe-S) clusters are not oxidized (53). Fnr activity is captured by three iModulons with similar activities in these samples; we therefore focus on Fnr-3, which has the highest explained variance of the three. ArcA and Fnr-3 activities are correlated (black line, **Figure 5B**) since they both sense different features of the same underlying cellular redox state.

**Figure 5.**
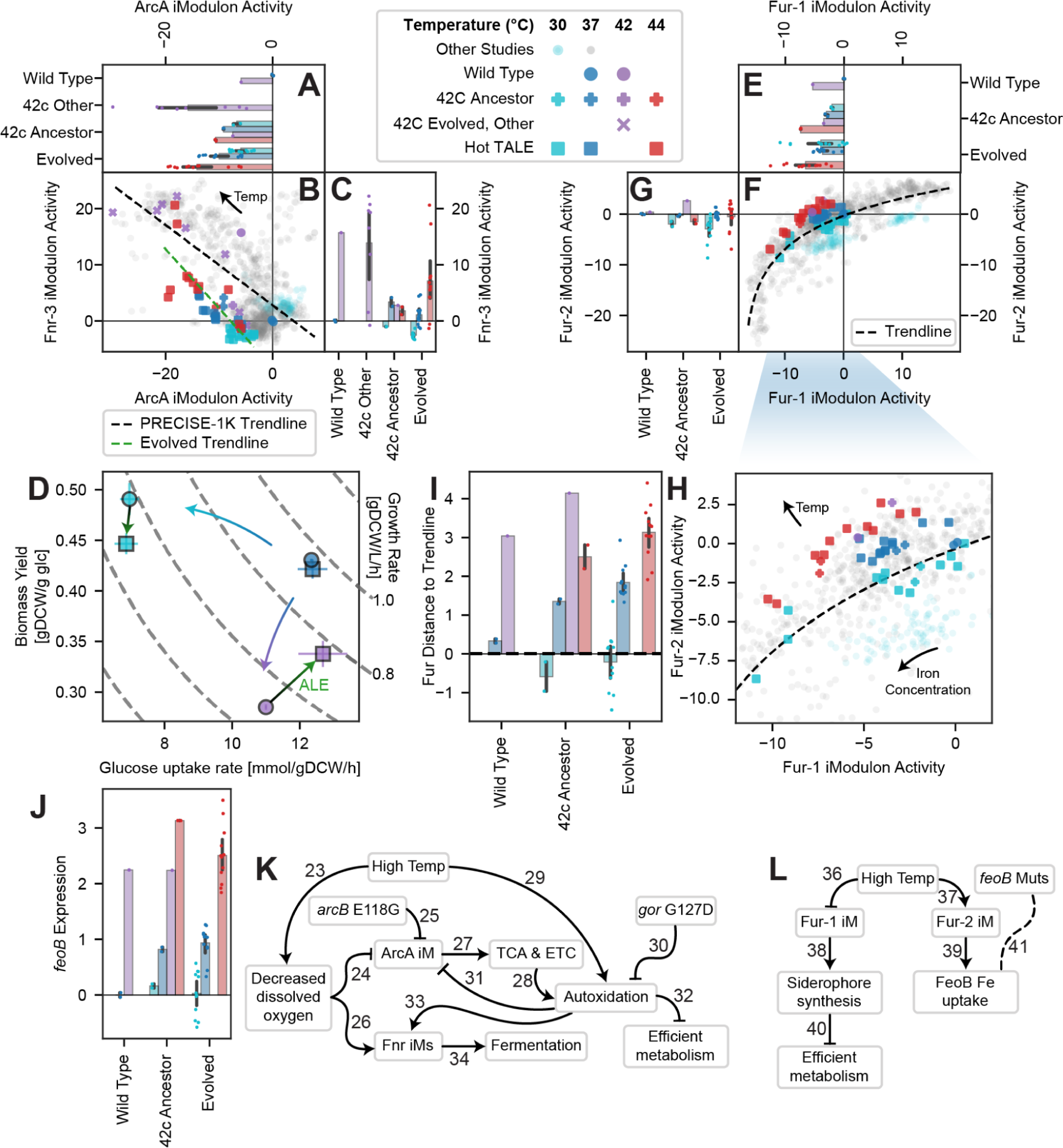
iModulon activities and mutations reveal hallmarks of redox metabolism, iron uptake, and uncharacterized genes which may facilitate temperature adaptation. All figures use the colors, and all scatterplots use the shapes, given in the legend (top middle). For bar graphs, error bars represent mean ± 95% confidence interval. **(A-C)** iModulon activities for ArcA, which regulates aerobic metabolism (by sensing oxidation state of quinones in the ETS), and Fnr-3, which regulates anaerobic metabolism (by sensing oxidative damage to the Fnr Fe-S cluster). In (B), linear fits for the evolved samples (green) and all other samples (black) are shown. Temperature shifts expression up the trendline toward a less oxidized state, and the evolved strains have shifted their trend leftward, likely due to an *arcB* E118G mutation. **(D)** Rate-yield plot demonstrating the effect of temperature and evolution on biomass yield and glucose uptake rate. Gray dotted lines indicate isoclines with constant growth rate according to the labels on the right. Colored arrows show the effect of a change in temperature (blue and purple) and evolution (green). **(E-G)** iModulon activities for Fur-1 and Fur-2, which regulate iron uptake and are fit to a logarithmic curve. **(H)** Zoomed in version of (F), showing that raising the temperature tends to shift samples above the trendline, toward Fur-2 expression. Fur-2 contains *feoABC*, the simple iron transporter, whereas Fur-1 contains the more metabolically expensive and less necessary siderophore synthesis pathways. The arrow labeled “Iron Concentration” shows the direction of increasing iron concentration from other studies in PRECISE-1K, and the one labeled “Temp” describes the observed trend from this study. **(I)** Distance to the trendline from panels (F) and (H), showing that increasing temperatures shifts the preference of Fur toward activating Fur-2. (J) *feoB* gene expression is correlated with temperature in all strains, which is consistent with the association between Fur-2 and high temperature. **(K-L)** Knowledge graphs for each of the temperature adaptation mechanisms presented in this figure, where numbering is consistent with Figure 2 and additional details are available in **Table S4**.

As temperature increases, gene expression shifts upward and leftward in the ArcA/Fnr-3 phase plane (**Figure 5A-C**). This shift indicates that high temperatures decrease oxidation, which is consistent with the decrease in oxygen solubility as temperatures increase (54). Indeed, a study described how higher temperatures can lead to oxygen limitation in animals (55). Decreased oxygen solubility may cause the ETC to be more reduced and the Fe-S clusters to be less oxidized, causing changes to the activity state of these iModulons. The expression change will tend to decrease the production of NADH and reliance on oxygen.

In addition to its effect on oxygen solubility, high temperature also increases the rate of autoxidation, a process in which reactive oxygen species (ROS) are generated by ETC components like NADH dehydrogenase (56, 57). Decreasing ArcA expression should decrease ETC activity and help to decrease the amount of electrons that end up being wasted by this process at high temperatures (56, 57). Interestingly, the 42c_3 strain and all fully evolved strains harbored the mutation *arcB* E118G, suggesting that they modified the ArcAB system to better tolerate high temperatures. In **Figure 5B**, we observe that ArcA iModulon activity has shifted to the left of the trendline formed by the other samples (8), so we infer that this mutation increases the phosphorylation of ArcA by ArcB (58). The *arcB* mutation would explain the shift in ArcA iModulon activities, and provide the benefit of decreasing autoxidation from the ETC, reinforcing a change induced by high temperatures. However, there ought to be a tradeoff to this mutation at lower temperatures, when autoxidation does not have as strong of an effect on biomass yield.

We also note one outlier strain, hot_9, which did not upregulate Fnr iModulons or further downregulate the ArcA iModulon when temperature increased. This strain harbored the mutation *gor* G127D, which may have enhanced ROS detoxification by glutathione reductase or decreased autoxidation (59) at high temperatures.

To explore the systems-level changes to energy metabolism that arise from these genomic and transcriptomic changes, we measured the glucose uptake rate and biomass yield of the wild type and evolved strains at three temperatures and plotted them on a rate-yield plane (**Figure 5D**). Regions of the rate-yield plane are associated with distinct states of energy metabolism called aero-types, as has been characterized in prior studies (13, 14, 60). Samples with high biomass yields are in the highest aero-type, corresponding to efficient aerobic growth. Lower aero-types are progressively less efficient and pump fewer protons across the inner membrane during respiration. Within an aero-type, samples may shift left or right based on the rate of glucose uptake. We find that temperature strongly affects yield and uptake: cold samples are highly efficient (high aero-type), but unable to rapidly uptake glucose (**Figure 5D**, blue arrow), whereas hot samples can rapidly take up glucose but have low yield (lower aero-type) due to heat-induced damage and waste (red arrow).

The evolved strains at high temperature have higher uptake and yield compared to the wild type (**Figure 5D**, green arrow labeled “ALE”). The increased uptake may be due to the changes toward anaerobic metabolism, which utilizes more glucose (51). The increased yield is the result of the combined success of many of the mutations which decrease temperature stress, including the decreased autoxidation brought about by the shift toward anaerobic metabolism. We note that, on average, the effects of evolution at 37°C are negligible. At 30°C, on the other hand (**Figure 5D**, green arrow with no label), yield decreases while glucose uptake rates remain low. This is consistent with the *arcB* mutation preventing upregulation of the high-yield aerobic pathways, which are highly efficient in the wild type at low temperatures.

Thus, iModulon and glucose uptake rate-yield analysis have revealed the effects of temperature and mutations on energy metabolism (**Figure 5K**). At high temperatures, dissolved oxygen decreases and electrons leak from the electron transport chain into ROS more readily (57), inducing a metabolic shift toward anaerobiosis which is amplified by an *arcB* mutation in the high temperature-tolerant evolved strains. A mutation in *gor* may alleviate some autoxidation at high temperatures. This shift successfully increases both glucose uptake and biomass yield at high temperatures, but carries a tradeoff that decreases yield at lower temperatures. This temperature tolerance strategy is informative for the fundamental biology of cross-stress tolerance and the relationship between stress and metabolism. Its mutations may also provide design variables of interest for fermentation applications which may experience high temperatures or uneven oxygenation.

### Fur preferentially derepresses *feoB*, a commonly mutated iron transporter

The two Fur iModulons regulate iron uptake systems and exhibit a nonlinear relationship (**Figure 5E-G**) (29). They explain approximately 1% of the variance in the transcriptome and exhibit an interesting relationship with temperature, in which temperature shifts activity perpendicularly to the trendline (**Figure 5H-I**). This behavior is observed in both wild type and evolved strains, suggesting that it may be a fundamental feature of fur binding. The effect is to prefer Fur-1 iModulon genes in lower temperatures and Fur-2 genes in higher temperatures. Fur-1 contains siderophore synthesis genes, which are needed when iron becomes less soluble at cold temperatures, as has been studied in *Vibrio salmonicida (61)*. Fur-2, on the other hand, contains less metabolically expensive ionic iron transporters, like *feoABC* (62), which is upregulated (**Figure 5J**). These would be preferred at higher temperatures due to their lower cost and the readily available dissolved iron (**Figure 5L**).

Though there are no transcriptional regulatory mutations to the iron uptake system, the transporter gene *feoB* mutates in three of the six evolved strains (hot_4: F363L; hot_8: W699*; hot_9: F363L & V563M). Further research ought to probe the effects of these mutations on the temperature stability and function of FeoABC.

### The *yjfIJKL* operon may be a new heat tolerance operon

Finally, a large 2.2% of the explained variance in the transcriptome is attributed to a single operon of all uncharacterized genes, *yjfIJKL*, which constitutes the YjfJ iModulon. The iModulon was named as such because *yjfJ* encodes what was previously thought to be a putative transcription factor, and it was presumed to be the regulator of the operon during the initial PRECISE-1K curation. The iModulon is strongly upregulated in 42c_3 and the evolved strains (**Figure 6A**), but not in any other samples in the dataset. This is predicted to be the result of a single nucleotide deletion 80 base pairs upstream of the operon (*yjfI-*pΔ1), in its promoter region (**Figure 6B**).

**Figure 6.**
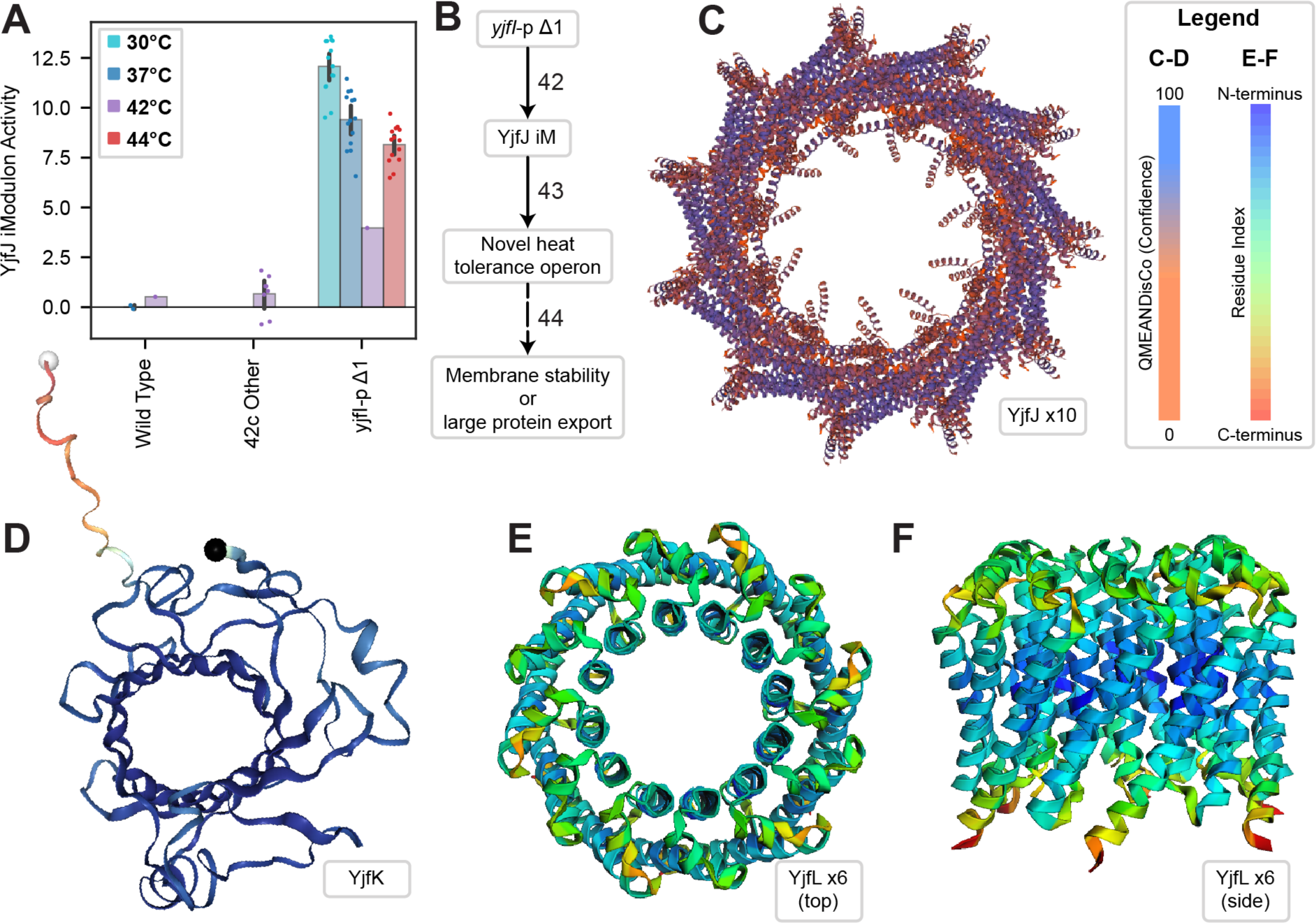
Upregulation and structure prediction for *yfjIJKL* suggest a role in membrane stability at high temperatures. **(A)** YjfJ iModulon activity, describing the expression of the *yjfIJKL* operon. The iModulon appears to be activated in all evolved strains by a single nucleotide promoter deletion upstream of *yjfI*. This iModulon represents an unknown molecular process, but is a clear signal detected by ICA. **(B)** Knowledge graph for this section, where numbering is consistent with Figure 2 and additional details are available in **Table S4**. **(C-F)** Predicted structures from AlphaFold (63) for YjfJ iModulon member genes. See **Table 1** for details. **(C)** YjfJ 55-mer colored by QMEANDisCo confidence score (64)). (**D)** YjfK 1-mer, also colored by QMEANDisCo. **(E-F)** YjfL 6-mer seen from top **(E)** and side view **(F)**. Colored by residue index as shown in legend.

**Table 1.** Structural predictions for uncharacterized genes in the strongly upregulated YjfJ iModulon. ESCRT-III: endosomal protein complex required for transport-III; QSPRD: Quaternary structure score (73); GMQE: Global model quality estimation (74); QMEANDisCo: Qualitative Model Energy Analysis with Distance Constraints (global) (64); pLLDT: Per-residue local distance difference test (75); iPTM: integrated predicted template modeling (76), PTM: predicted template modeling (63); Model: 0.8*iPTM + 0.2*PTM (77).

| Gene | Description | Similar To | Location | Structure | Confidence |
| --- | --- | --- | --- | --- | --- |
| **Yjfl**<br>**(b4181)** | Alpha helix and beta sheet domains, low confidence | Membrane lipo-proteins that form periplasmic sheets | Periplasm | N/A | N/A |
| **YjfJ**<br>**(b4182)** | **Figure 6D.** 55-mer pore structure with C-11 symmetry | PspA, Vipp1, and other ESCRT-III proteins that remodel membranes | Inner membrane | SWISS-MODEL (6zvr) | QSPRD: 0.22<br>GMQE: 0.67<br>QMEANDisCo: 0.63<br>Seq: 20.64% |
| **YjfK**<br>**(b4183)** | **Figure 6E.** Primitive outer membrane B-barrel porin | MliC, YqeJ, YnfC | Outer membrane | AlphaFold | pLDDT: 0.925 |
| **YjfL**<br>**(b4184)** | **Figure 6F-G.** Multiple transmembrane helices. Hexamer forms inner membrane channels. | Channel-like porins | Inner membrane | AlphaFold Multimer (unrelaxed) | iPTM: 0.67<br>PTM: 0.71<br>Model: 0.68 |

Given the large influence on the transcriptome, the likely causal mutation, and the lack of existing knowledge about these genes, we decided to perform structure prediction in order to improve functional annotation of the YjfJ iModulon and identify its possible role in high temperature stress mitigation. Prior to this work, it was known that YjfJ exhibits homology to PspA, a phage shock protein (65), which we confirmed using ssbio (41.8% similarity, score 88.5) (66). PspA uses a variety of interesting structural mechanisms to remodel bacterial membranes (67, 68). Using QSPACE (69), we were able to quickly find all structures for YjfJ in the SWISS-MODEL repository (70), AlphaFold Database (63, 71), and Protein Data Bank (PDB) (72). The SWISS-MODEL structure for YjfJ resembles an ESCRT-III-like membrane channel (**Figure 6D**) (68), suggesting that YjfJ can also play a role in membrane remodeling, repair, and maintenance. We also performed structural prediction for the other three genes in the iModulon: YjfK is likely a primitive outer membrane porin (**Figure 6E**), YjfL may oligomerize into inner membrane channel-like porin structures (**Figure 6F-G**), and YjfI had only low-confidence structure predictions typical for relaxed membrane lipo-proteins that can form sheet-like structures in the periplasm. Details of each gene are summarized in **Table 1**.

Taken together, all four genes in the iModulon likely interact with the cellular envelope, and three of them (*yjfJKL*) appear to add pore structures to the membranes. Channels like these may improve membrane stability in high temperature conditions by exerting lateral pressure across the phospholipid bilayer, packing the phospholipids together despite local instability within the holes (78). There is also a chance that larger openings formed by YjfJ are able to secrete misfolded proteins from the periplasm. Thus, the mechanisms associated with the YjfJ iModulon are similar to those for the motility and fimbriae iModulons discussed earlier: the evolved cells upregulate certain envelope-associated structures, with predicted effects that increase membrane stability or protein aggregate efflux. The discovery of the YjfJ iModulon, its upregulation during temperature adaptation, and its putative functional annotation through structural proteomics, provide clear impetus for undertaking a future study to detail is molecular functions. The promoter mutation we identified would be useful to increase expression in such a study.

## Conclusion

In this study, we used ALE to produce six *E. coli* strains which can grow at 45.3°C, a temperature lethal to wild type cells. Though their hypermutator phenotype made a detailed global analysis of their genomic changes intractable, important features of the strains’ tolerance strategies were revealed via an iModulon analysis of their transcriptome. We discussed mechanisms that involve 11 mutations and 14 iModulons, that represent about half of all variation in the gene expression observed in these strains (**Figure 2**). The strains use gene expression adaptations to improve temperature tolerance by (i) specializing their stress response by downregulating RpoS and upregulating RpoH, (ii) activating flagellar basal bodies and fimbriae while downregulating FliA with possible effects on envelope stability or the export of heat-damaged, misfolded proteins, (iii) downregulating aerobic metabolism genes to counteract changes to oxygen solubility and autoxidation rates, (iv) upregulating and modifying ionic iron uptake while shifting away from unnecessary expression of siderophores, and (v) upregulating the previously uncharacterized *yjfIJKL* operon, which based on our structural analysis is likely to improve membrane stability.

The five mechanisms described above suggest three general principles for mesophilic microbes growing at high temperatures. The first is to streamline stress responses and metabolism – the strains downregulate the RpoS general stress response and the autoxidation-inducing ArcA regulon. The shift from Fur-1 to Fur-2 also helps to streamline metabolism by de-emphasizing siderophore pathways, though this may be a wild type phenomenon and not an evolved feature. Secondly, the strains need to deal with protein aggregation that occurs at high temperatures. To do so, they can upregulate the RpoH sigmulon of proteases and chaperones (79), and they may also use the flagellar basal bodies and fimbriae machinery to export misfolded proteins. Thirdly, the strains improve envelope stability by upregulating membrane protein systems, including the flagellar basal body, fimbriae, and *yjfIJKL* operon.

This study and similar work on ROS tolerance (14) emphasize the value of transcriptomic analysis through iModulons for building a multi-level understanding of cellular stress tolerance phenotypes. In both cases, the stress response becomes specialized for the given strain by modifying activity of RpoS while leaving the specific stress regulon (RpoH or SoxS) to function as it does in wild type. Interestingly, both cases also showed a shift toward anaerobiosis and higher preference for Fur-2 ionic iron transport, but with different predicted underlying mechanisms. The rich information gleaned from these ALE experiments and transcriptomic datasets motivates further applications of iModulons for understanding unique strains. This effort will build up more examples associated with each iModulon and further enrich the field’s working understanding of transcriptional regulation.

A particularly fruitful use of iModulon analysis in this study lies in the use activity phase planes (**Figures 3B, 4D-E, 5B, 5F, 5H)**. Each figure showed a trend that was observed across the >1000 samples of PRECISE-1K (8), along with modifications to the overall trend resulting from regulatory changes in the evolved strains. This is an example of an emerging principle of data science in microbial physiology – *i.e.* learning with scale – as the evolved transcriptomic changes could not have been understood without the context of a large reference dataset.

The goal of this study was to identify mechanisms underlying temperature tolerization. iModulons proved to be a key tool for achieving this goal, particularly because mutational data was complicated due to hypermutator phenotypes. Relating between iModulons and selected mutations is important, but we recognize that this pursuit is somewhat limited in scope. The mutational mechanisms are inferred based on literature associations of the genes and regulons, as opposed to being individually experimentally validated. We rely on prior work in the literature, which allows us to cover more of the global features of the transcriptome in a single manuscript. This approach bears the risk of presenting incorrect conclusions, and thus we encourage future studies to more thoroughly validate the hypotheses presented here using traditional methods.

In addition to its contribution to the understanding of TRN adjustments at high temperatures, we anticipate that this study will lead to practical applications. Engineering flagellar basal bodies for heterologous protein export is a promising approach (44), and we have implicated mutations in the ATPase genes *fliIJ* in a mechanism that upregulates the export basal body without the wasteful production of other motility proteins. We are also the first to report that high temperatures change the activity of ArcA and Fnr and predict that it is due to their sensitivity to temperature-dependent changes in oxygen. This observation could also be useful for designing cell factories, in which changes to oxygen and temperature commonly occur, and regulatory effects need to be precisely understood. Also, temperature-tolerant pathogenic *E. coli* strains would be more able to survive fever conditions in a host, so the mutations and mechanisms described here could help to explain the evolution of pathogenic strains.

Taken together, we presented a global characterization of laboratory evolved, high temperature-tolerant strains of *E. coli* with emphasis on the transcriptome as opposed to the genome. Our multi-level approach was effective for understanding the coordination of multiple mechanisms resulting in temperature tolerization. It also predicted new mechanisms involved in temperature tolerization and resulted in the putative functional annotation of unknown genes. Given the availability of large amounts of transcriptomic data and tools like iModulon analysis, we believe that TRN evolution will continue to be elucidated for a variety of environmental challenges. Such studies will reveal how known cellular processes cooperate in generating tolerized phenotypes, and will discover new ones.

## Materials & methods

### Resource availability

RNA-seq data have been deposited to GEO and are publicly available as of the date of publication, under accession number GSE140478. DNA-seq data are available from aledb.org under the project “Hot mutL”. iModulons and related data are available from iModulonDB.org under the dataset “*E. coli* PRECISE-1K”. The YjfJ 55-mer structure is available at https://swissmodel.expasy.org/repository/uniprot/P0AF78?template=6zvr.1.A&range=1-231.

All original code and data to generate figures are available at github.com/SBRG/Hot-ALE, which also links to the alignment, ICA, and iModulon analysis workflows (80). It has been deposited at Zenodo and is publicly available as of the date of publication.

Any additional information required to reanalyze the data reported in this paper, including strains generated in this study, is available from the lead contact upon request: Bernhard Palsson.

### Microbial strains

The starting strain of the original 42C evolution(Sandberg et al. 2014) was *E. coli* K-12 MG1655. Mutations for the evolved strains are listed on aledb.org and in **Table S1**.

### Culture conditions

All strains were grown and evolved in M9 minimal medium prepared by addition of 0.1 mM CaCl_2_, 2 mM MgSO_4_, 1x trace elements solution, 1x M9 salt solution, and 4 g/L D-glucose to Milli-Q water. The M9 salt solution was composed of 68 g/L Na_2_HPO_4_, 30 g/L KH_2_PO_4_, 5 g/L NaCl, and 10 g/L NH_4_Cl. The trace elements solution was prepared by mixing 27 g/L FeCl_3_・6 H_2_O, 1.3 g/L ZnCl_2_, 2 g/L CoCl_2_ ⋅ 6 H_2_O, 2 g/L Na_2_MoO_4_⋅ 2 H_2_O, 0.75 g/L CaCl_2_, 0.91 g/L CuCl_2_, and 0.5 g/L H_3_BO_3_ in a Milli-Q water solution consisting of 10% concentrated HCl by final volume. Sterilization was achieved in all solutions and media by filtration through a 0.22 μM PVDF membrane.

### Adaptive laboratory evolution

Stage I of the ALE experiment was started from isolates of the wild-type E. coli K-12 MG1655, and evolved at 42°C as described previously (22). Clones were isolated from ten populations at the end of this experiment, and eight of them with distinct mutational histories were used to start the Stage II ALE experiment. Unfortunately, a contamination event early in the Stage II ALE led to the 42c_3 strain becoming the dominant strain in all flasks that were subsequently analyzed, as evidenced by DNA resequencing. Although this event led to less diverse starting conditions than were originally intended, it does suggest that mutations in 42c_3 were particularly beneficial in the ALE conditions, and diverse endpoint strains were still obtained.

All cultures during the Stage II evolution were grown in 35 mL flasks with a 15 mL working volume, and were vigorously stirred at 1100 rpm to create a well-mixed and aerobic environment. Initial temperatures for these cultures were set to 42°C. The temperatures were increased by 0.5°C approximately every 150 generations (∼15 passages) to give the cultures time to optimize their growth under the new conditions. Due to the higher stress levels, temperature increases were only 0.25°C above 44°C. An automated system was used to propagate the evolving populations over the course of the ALE. To maintain the evolving population at the exponential growth phase, their growth was periodically monitored by taking optical density measurements at a 600 nm wavelength (OD600) on a Tecan Sunrise reader plate (**Figure S1**). Once reaching the target OD600∼0.3 (∼1 on a 1 cm path length spectrophotometer), approximately 0.66% of the cells in a population were passaged to the fresh medium. Population samples along the adaptive trajectories were taken by mixing 800 μL of culture with 800 μL of 50% glycerol, and stored at -80°C for subsequent analysis (not reported).

### DNA sequencing and mutation calling

Growth-improved clones along the ALE trajectory were isolated and grown in the standard medium condition. Cells were then harvested while in exponential growth and genomic DNA was extracted using a KingFisher Flex Purification system previously validated for the high throughput platform mentioned below (81). Shotgun metagenomic sequencing libraries were prepared using a miniaturized version of the Kapa HyperPlus Illumina-compatible library prep kit (Kapa Biosystems). DNA extracts were normalized to 5 ng total input per sample using an Echo 550 acoustic liquid handling robot (Labcyte Inc), and 1/10 scale enzymatic fragmentation, end-repair, and adapter-ligation reactions carried out using a Mosquito HTS liquid-handling robot (TTP Labtech Inc). Sequencing adapters were based on the iTru protocol (82), in which short universal adapter stubs are ligated first and then sample-specific barcoded sequences added in a subsequent PCR step. Amplified and barcoded libraries were then quantified using a PicoGreen assay and pooled in approximately equimolar ratios before being sequenced on an Illumina HiSeq 4000 instrument.

Sequencing reads were filtered and trimmed using AfterQC version 0.9.7 (83). We mapped reads to the *E. coli* K-12 MG1655 reference genome (NC_00913.3) using the breseq pipeline version 0.33.1 (84). Mutation analysis was performed using ALEdb (4).

### Physiological characterization

Cultures were initially inoculated from -80°C glycerol stocks, and grown at 37°C overnight. Physiological adaptation was achieved by growing cell cultures exponentially over 2 passages for 5 to 10 generations at the target temperature for phenotypic characterization. Next, cultures growing at the exponential growth phase were passaged to a 15 mL working volume tube and grown fully aerated. Spectrophotometer readings at OD600 were periodically taken (Thermo Fisher Scientific, Waltham, MA) until stationary phase was reached. Growth rates were determined for each culture by least-squares linear regression of ln(OD600) versus time.

Samples were filtered through a 0.22 micrometer filter (MilliporeSigma, Burlington, MA) at the same time OD600 measurements were taken, and the filtrate was analyzed for glucose and acetate concentrations using a high-performance liquid chromatography system (Agilent Technologies, Santa Clara, CA) with an Aminex HPX-87H column (Bio-Rad Laboratories, Hercules, CA). Glucose uptake rates and acetate production rates in exponential growth were determined by best-fit linear regression of glucose and acetate concentrations versus cell dry weights, multiplied by growth rates over the same sample range. The above described phenotypic characterizations were performed for two biological replicates of each of the selected clonal isolates along the ALE trajectory, at 30°C, 37°C, and 44°C, respectively.

### RNA sequencing

During phenotypic characterization, 3 mL of cell broth was taken at OD600∼0.6, and immediately added to 2 volumes of Qiagen RNAprotect Bacteria Reagent (6 mL). Then, the sample was vortexed for 5 seconds, incubated at room temperature for 5 minutes, and immediately centrifuged for 10 minutes at 5000g. The supernatant was decanted, and the cell pellet was stored in the -80°C. Cell pellets were thawed and incubated with Readylyse Lysozyme, SuperaseIn, Protease K, and 20% SDS for 20 minutes at 37°C. Total RNA was isolated and purified using the RNeasy Plus Mini Kit (Qiagen) columns following vendor procedures. An on-column DNase-treatment was performed for 30 minutes at room temperature. RNA was quantified using a Nanodrop and quality assessed by running an RNA-nano chip on a bioanalyzer. The rRNA was removed using Illumina Ribo-Zero rRNA Removal Kit (Gram-Negative Bacteria). The quantity was determined by Nanodrop 1000 spectrophotometer (Thermo Scientific). The quality was checked using RNA 6000 Pico Kit using Agilent 2100 Bioanalyzer (Agilent). Paired-end, strand-specific RNA-seq library was built with the KAPA RNA Hyper Prep kit (Kapa Biosystems) following manufacturer’s instructions. Libraries were sequenced on an Illumina HiSeq 4000 instrument.

As part of the PRECISE-1K dataset (8), transcriptomic reads were mapped using our pipeline (https://github.com/avsastry/modulome-workflow) (80) and run on Amazon Web Services Batch. First, raw read trimming was performed using Trim Galore with default options, followed by FastQC on the trimmed reads. Next, reads were aligned to the *E. coli* K-12 MG1655 reference genome (NC_000913.3) using Bowtie (85). The read direction was inferred using RSeQC (86). Read counts were generated using featureCounts (87). All quality control metrics were compiled using MultiQC (88). Finally, the expression dataset was reported in units of log-transformed transcripts per million (log(TPM)).

All included samples passed rigorous quality control, with “high-quality” defined as (i) passing the following FastQC checks: *per_base_sequence_quality, per_sequence_quality_scores, per_base_n_content, adaptor content;* (ii) having at least 500,000 reads mapped to the coding sequences of the reference genome (NC_000913.3); (iii) not being an outlier in a hierarchical clustering based on pairwise Pearson correlation between all samples in PRECISE-1K; and (iv) having a minimum Pearson correlation between biological replicates of 0.95.

### iModulon computation and curation

The full PRECISE-1K compendium, including the samples for this study, was used to compute iModulons using our previously described method (8, 89). The log(TPM) dataset **X** was first centered such that wild-type *E. coli* MG1655 samples in M9 minimal media with glucose had expression values of 0 for all genes. Independent component analysis was performed using the Scikit-Learn (v0.19.0) implementation of FastICA (90). We performed 100 iterations of the algorithm across a range of dimensionalities, and for each dimensionality we pooled and clustered the components with DBSCAN to find robust components which appeared in more than 50 of the iterations. If the dimensionality parameter is too high, ICA will begin to return single gene components; if it is too low, the components will be too dense to represent biological signals. Therefore, we selected a dimensionality which was as high as possible without creating many single gene components, as described (89). At the optimal dimensionality, the total number of iModulons was 201. The output is composed of matrices **M** [genes x iModulons], which defines the relationship between each iModulon and each gene, and **A** [iModulons x samples], which contains the activity levels for each iModulon in each sample.

For each iModulon, a threshold must be drawn in the **M** matrix to determine which genes are members of each iModulon. These thresholds are based on the distribution of gene weights. The highest weighted genes were progressively removed until the remaining weights had a D’agostino K^2^ normality below 550. Thus, the iModulon member genes are outliers from an otherwise normal distribution. iModulon annotation and curation was performed by comparing them against the known TRN from RegulonDB (91). Names, descriptions, and statistics for each iModulon are available from the PRECISE-1K manuscript (8), iModulonDB (9), and **Table S3.**

### Differential iModulon activity analysis

DiMAs were calculated as previously described (6, 80). For each iModulon, a null distribution was generated by calculating the absolute difference between activity levels in each pair of biological replicates and fitting a log-normal distribution to them. For the groups being compared, their mean difference for each iModulon was compared to that iModulon’s null distribution to obtain a p-value. The set of p-values for all iModulons was then false discovery rate (FDR) corrected to generate q-values. Activities were considered significant if they passed an absolute difference threshold of 5 and an FDR of 0.1. The main comparison in this study was between the wild type strain at 42°C (n = 1) and the combined set of all fully evolved strains at 44°C (n = 12). This comparison is shown in **Figure 1F**, and its p-values are reported in figure captions throughout the manuscript as well as in **Table S3**. We used the same statistical algorithm to compare the evolved strains at 30°C (n = 12) and 44°C (n = 12) in **Figure 1G**.

### iModulon explained variance calculation

The explained variance for each iModulon in this study was calculated using our workflow (80). Since iModulons are built on a matrix decomposition, the contribution of each one to the overall expression dataset can be calculated. For each iModulon, the column of **M** and the row of **A** for the evolved samples in this study were multiplied together, and the explained variance between the result and the full expression dataset was computed. These explained variance scores were used to size the subsets of the treemap in **Figure 1D** and the icons in the third column of **Figure 2**. Note that the variance explained by ICA is ‘knowledge-based’ in contrast to the ‘statistic-based’ variance explanation provided by the commonly used principal component analysis (PCA).

### Structure Comparison & Prediction

The QSPACE platform (69) was used to identify protein structures for YjfJ (b4182), YjfK (b4183) and YjfL (b4184) available in the PDB (72), the SWISS-MODEL repository (70), and the AlphaFold Database (63, 71). The SWISS-MODEL for YjfJ (P0AF78-6zvr.1.A) was derived from a template of the ESCRT-III-like protein Vipp1 with C-11 symmetry (PDB:6ZVR, (92)), however C12-18 symmetries have also been proposed for Vipp1((92) and (93)). Vipp1 is homologous to PspA (92). A high-confidence (pLDDT = 0.925) AlphaFold model (P39293) was identified for YjfK in the AlphaFold Database. Although the shape of the YjfK AF-model resembles a small outer membrane channel, neither the sequence-based prediction (DeepTMHMM (94)) nor structure-based prediction (OPM PPM server 2.0 (95)) were able to confirm that YjfK belongs to the membrane. The YjfL hexamer was modeled using ColabFold (96) (AF-Multimer v2, model score = 0.68).

## Acknowledgements

We thank Amitesh Anand, Xin Fang, and Patrick V. Phaneuf for helpful discussions. This work was funded by National Institutes of Health grants (awards GM102098 and GM057089), and the Novo Nordisk Foundation (awards NNF10CC1016517 and NNF20CC0035580). This research used resources of the National Energy Research Scientific Computing Center, supported by the U.S. Department of Energy under Contract No. DE-AC02-05CH11231.

## Author contributions

K.R., K.C., A.M.F., and B.O.P. designed the study. K.C., C.A.O., T.E.S., Y.G., S.X., Y.H., and R.S. performed experiments. K.R., K.C., and E.A.C. analyzed the data and wrote the manuscript, with contributions from all co-authors.

## Appendix

### Supplemental Figures

**Figure S1.**
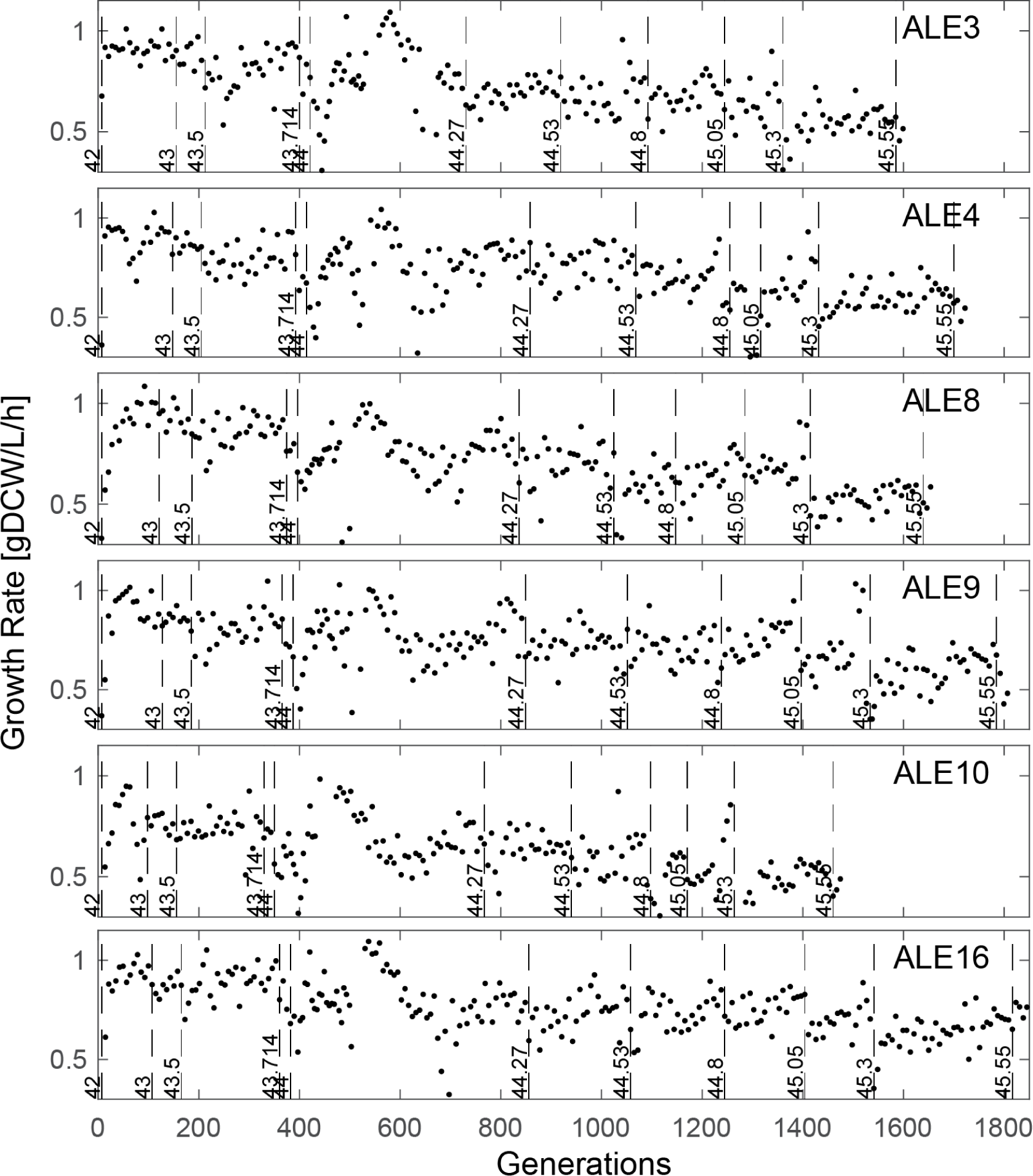
Growth rates and temperatures for each flask in ALE. Dotted vertical lines indicate the flasks at which the temperature was increased. Generation numbers are estimated from the growth rate and elapsed time of each flask.

### Supplemental Tables

***Table S1: Mutations in 42c_3 starting strain.***

Mutations details, position, type, sequence change, and affected genes were generated by the ALEdb mutation calling pipeline (4). Positions refer to locations in the NC_000913 genome (97)The ‘Figure’ column contains the figure number in which a mutation is discussed, with a * for *mutL* which is discussed for its relevance to the hypermutator phenotype but not in a particular figure. This is a subset of **Table S2** that only includes the mutations that were present in the starting strain.

***Table S2. Mutations in high temperature tolerized strains.***

Mutations details, position, type, sequence change, and affected genes were generated by the ALEdb mutation calling pipeline (4). ‘Category’ was used to generate **Figure 1C**. The ‘Figure’ column is blank for mutations that are not discussed in the paper, and references a figure number otherwise (* for *mutL*, which is discussed but not in a figure). Columns labeled with strain numbers indicate presence or absence of the mutation in the given strain.

***Table S3. iModulon activities in high temperature tolerized strains.***

iModulons are listed by explained variance in the evolved strains’ data. If the iModulon was significantly differentially activated in **Figure 1F**, ‘Evolution P’ indicates the false discovery rate corrected p-value and ‘Evolved Minus WT’ shows the difference in mean activities. Similarly, if the iModulon was significantly differentially activated in **Figure 1G**, then statistics are listed under ‘Temperature P’ and ‘44C Minus 30C’ where positive differences indicate upregulation at 44°C relative 30°C. ‘Category’ was used to generate **Figure 1D**. ‘Figure’ lists the figure that is relevant for understanding the iModulon’s behavior, if applicable. Remaining descriptive columns are copied from the PRECISE-1K curation of these iModulons (8). See iModulonDB.org for details of each iModulon, including its member genes, activity levels across over 1000 conditions including those from this study, and overlap with associated regulons.

***Table S4. Meaning, evidence, and novelty for each proposed connection***

Each row of this table corresponds to an arrow in **Figure 2**, as well as **3D, 4C, 5K-L,** and **6B**. The ‘From’, ‘To’, and ‘Meaning’ columns describe the relationship in more detail than can be displayed on a knowledge graph. ‘Figure’ contains the specific relevant figure panels if available. ‘Data Evidence’ contains written descriptions of the evidence from this study associated with the relationship. ‘Literature Evidence’ lists citations which agree with or establish the relationship. The ‘New?’ column assesses the novelty of the relationship, which may be previously established, new in this study, or describe the new details and perspectives revealed by comparing the literature with the new data.

### Note S1. Flagellar iModulons provide additional insights into the regulation of motility, related to Figure 4

The two iModulons nearly perfectly mirror the known regulation (**Figure 4B**). However, FlhDC-2, but not FliA, includes *fliLMNOPQR*, despite the fact that it has binding sites for both regulators. This operon is needed earlier in the synthesis of flagella (46). iModulons learn from relationships in expression data, so the exclusion of this operon from FliA indicates that these genes tend to be more correlated with the early stages of synthesis, in agreement with their function and despite the ability to be regulated by the late stage regulator. In addition, a third iModulon plays a unique role: the FlhDC-1 iModulon contains several genes from both iModulons at both positive and negative weights, and appears to capture a third dimension or nonlinearity in the motility transcriptome. This may reflect different binding affinities and changing ratios between the two regulators. FlhDC-1 and 2 form a nonlinear curve, suggesting that samples adjust towards FlhDC-1 as FlhDC expression increases (**Figure 4E**). Thus, iModulons reflect the known transcriptional regulation while providing additional nuance which is useful for a practical understanding of the system.

